# Systematic benchmark of state-of-the-art variant calling pipelines identifies major factors affecting accuracy of coding sequence variant discovery

**DOI:** 10.1101/2021.04.13.439626

**Authors:** Ruslan Abasov, Varvara E. Tvorogova, Andrey S. Glotov, Alexander V. Predeus, Yury A. Barbitoff

## Abstract

Accurate variant detection in the coding regions of the human genome is a key requirement for molecular diagnostics of Mendelian disorders. Efficiency of variant discovery from nextgeneration sequencing (NGS) data depends on multiple factors, including reproducible coverage biases of NGS methods and the performance of read alignment and variant calling software. In this work, we systematically evaluated the performance of 4 popular short read aligners (Bowtie2, BWA, Isaac, and Novoalign) and 6 variant calling and filtering methods (based on DeepVariant, Genome Analysis ToolKit (GATK), FreeBayes, and Strelka2) using a set of 10 “gold standard” WES and WGS datasets. Our results suggest that Bowtie2 performs significantly worse than other aligners and should not be used for medical variant calling. When other aligners were used, the accuracy of variant discovery mostly depended on the variant caller and not the read aligner. Among the tested variant callers, DeepVariant consistently showed the best performance and the highest robustness, with other state-of-the-art tools, i.e. Strelka2 and GATK with 1D convolutional neural network variant scoring, also showing high performance on both WES and WGS data. The results show surprisingly large differences in the performance of cutting-edge tools even on high confidence regions of the coding genome. This highlights the importance of regular benchmarking of quickly evolving tools and pipelines. Finally, we discuss the need for a more diverse set of gold standard genomes (e.g. of African, Hispanic, or mixed ancestry) that would allow to control for deep learning model overfitting. For similar reasons there is a need for better variant caller assessment in the repetitive regions of the coding genome.

## Introduction

Over the past decade next-generation sequencing (NGS) has become a widely used technique in genetics and genomics (van Dijk et al., 2014). Rapid technology development, as well as the introduction of whole-exome sequencing (WES) and target gene sequencing panels, facilitated NGS application to the analysis of human genome variation and molecular diagnostics of inherited disease. Millions of individual exomes and genomes have been sequenced across the globe, and large-scale variant datasets have been constructed from NGS data, including the Genome Aggregation Database (gnomAD) (Karczewski et al., 2020) and UK Biobank exome sequencing dataset (Bycroft et al., 2016). These datasets are extensively used in both clinical practice and basic human genetics research.

Despite huge developments over the past years, accuracy and reliability of variant discovery (variant calling) from NGS data still has room for improvement. Any basic variant calling pipeline includes two key stages: read alignment against a reference genome sequence and variant calling itself. Hence, quality of the reference genome sequence (Barbitoff et al., 2018) as well as properties of the software tools used for read alignment and variant calling all influence the final result. While BWA is considered a gold standard solution for short read alignment in medical genetics (van der Auwera et al., 2013), several other aligners have been developed and are commonly used, including Bowtie2 (Langmead et al., 2012), Isaac (Illumina Inc. USA), and Novoalign (Novocraft Technologies, USA). The spectrum of software tools for variant calling is much broader, ranging from relatively simple (such as the SAMtools/BCFtools pipeline (Li et al., 2009)) to rather complex ones (e.g., Genome Analysis ToolKit (GATK) (McKenna et al., 2010; DePristo et al., 2011; van der Auwera et al., 2013) HaplotypeCaller based on local haplotype assembly and Markov model-based genotyping). Active development of deep learning models and their application to biological data led to the introduction of neural network-based variant discovery methods such as DeepVariant (Poplin et al., 2018). Variant filtration methods based on convolutional neural networks are now also available in the most recent versions of GATK.

Existence of multiple variant calling pipelines predicates the need for a gold standard genome variation dataset that can be used for extensive benchmarking of variant discovery pipelines. Such gold standard dataset has been compiled by the Genome In A Bottle Consortium (GIAB) and the National Institute of Standards (NIST) (Zook et al., 2016). The dataset includes high-confidence genotypes for a set of samples (the European NA12878/NA12891/NA12892 trio, the Chinese trio, and the Ashkenazi trio) obtained using multiple genotyping strategies. These high-confidence variant calls can be used as a truth set to evaluate the accuracy of variant calling, and estimate the precision and sensitivity of variant discovery. The GIAB gold standard dataset has been used multiple times for benchmarking of variant detection solutions. For example, a 2015 study by Hwang et al. (Hwang et al., 2015) used a set of sequencing datasets of the NA12878 sample and demonstrated important differences in the accuracy of variant calling pipelines available at the time, with a combination of BWA-MEM and SAMtools being the best solution for SNP calling, and BWA-MEM and GATK-HC for indels. More recently, several comparative analyses have shown that DeepVariant and Strelka2 (Kim et al., 2018) show the best performance on individual GIAB samples (Supernat et al., 2018; Chen et al., 2019; Zhao et al., 2020). The most recent comparative evaluation also demonstrated the utility of combining variant calling results from several pipelines (Zhao et al., 2020). While the aforementioned studies provide important information regarding the performance of different software, a single gold standard sample (NA12878) is usually used for comparison. This limitation does not allow estimation of the robustness of different pipelines and their ability to call variants in samples of different origin and/or sequencing quality.

In 2019, best practices for benchmarking variant calling software have been developed by the Genome Alliance for Genomics and Health (GA4GH) (Krusche et al., 2019). A reference implementation of the GA4GH benchmarking strategy, hap.py, allows researchers to evaluate the performance of a variant calling pipeline in a stratified manner, i.e. compare the accuracy of variant discovery in different sets of regions and for different variant types (Krusche et al., 2019). Such a stratified approach provides important information regarding the major factors affecting variant discovery. This, in turn, gives an opportunity to conduct a systematic survey of factors affecting reliable variant discovery. Previously, we have conducted a detailed analysis of the determinants of human coding sequence coverage in WES and WGS (Barbitoff et al., 2020). This study showed that all modern approaches to human genome resequencing have reproducible coverage bias, and mappability limitations are its major driver. These results prompted us to investigate the influence of different sequence-based factors and coverage biases on the performance of variant calling software. To this end, in this work we applied 24 different combinations of read alignment, variant calling, and variant filtration tools to a set of 10 gold standard samples from the GIAB data, and evaluated their robustness and general performance across different sets of human coding sequences.

## Materials and Methods

### Data acquisition

For this analysis, five GIAB samples were selected: the NA12878 (HG001), three members of the Ashkenazi trio (HG002, HG003, and HG004), and one representative of the Chinese trio (HG005). Gold standard data for these samples were downloaded from the GIAB FTP repository (all WGS samples) or the NCBI Sequencing Read Archive SRA (all WES samples, respective SRA IDs: ERR1905890, SRR2962669, SRR2962692, SRR2962694, and SRR2962693). All sequencing datasets were generated using the Illumina Hiseq platform (HiSeq 2500 for all WGS datasets and HG002-HG005 WES samples; HiSeq 4000 for HG001 WES). Truth variant sets and high-confidence variant calling regions in BED format (release v. 3.3.2) were downloaded from the GIAB FTP data repository (https://ftp-trace.ncbi.nlm.nih.gov/giab/ftp/release/). High-confidence regions for the HG001 sample were broader compared to HG002-HG005 (31.7 Mbp of coding sequences were included into the high-confidence regions for HG001, and only 29.2 - 29.9 Mbp for HG002-HG005). Reference stratification BED files were retrieved from the GitHub repository provided by GIAB (https://github.com/genome-in-a-bottle/genome-stratifications/blob/master/GRCh37/v2.0-GRCh37-stratifications.tsv).

### Variant calling pipelines

Variant discovery and filtering was performed using 24 different strategies, with 4 different read alignment software and 4 modern variant calling solutions used. For read alignment, we used BWA MEM v.0.7.17 (Li and Durbin, 2009); Bowtie2 (Langmead et al., 2012), Novoalign v. 4.02.01 (http://novocraft.com/novoalign/) and Isaac v. 04.18.11.09 (https://github.com/Illumina/Isaac4). Novoalign was used under trial license obtained by R.X.A. and Y.A.B. Reads were aligned against the GRCh37.p13 primary human reference genome sequence. Aligned reads were pre-processed using GATK to mark duplicate read pairs. Coverage statistics were collected using GATK v. 4.1.4.1 (McKenna et al., 2010). GENCODE v19 exon coordinates were used to evaluate the depth and breadth of coverage. Coverage of the high-confidence CDS regions in all samples was analyzed using the results of read alignment with BWA MEM v. 0.7.17.

Variant calling was performed using four different tools: FreeBayes v. 1.3.1 (Garrison and Marth, 2012), GATK HaplotypeCaller (HC) v. 4.1.4.1 (McKenna et al., 2010; DePristo et al., 2011), Strelka2 v. 2.9.10 (Kim et al., 2018), and DeepVariant v. 0.10.0 (Poplin et al., 2018). The DeepVariant caller was used with the default model for WGS or WES data, respectively. For the GATK HaplotypeCaller, deduplicated reads in BAM format were also preprocessed using base quality score recalibration according to GATK Best Practices (https://gatk.broadinstitute.org/hc/en-us/articles/360035535932-Germline-short-variant-discovery-SNPs-Indels-). Variants were called in a single-sample mode, and the resulting VCF was subject to variant filtration using CNNScoreVariants with different model types (reference-based (1D) or reads-based (2D)) and hard filtering with the recommended parameters. For both CNN models, different tranche values were tested, and SNP tranche value of 99.9 and indel tranche value of 99.5 were used as showing the best performance. For CNN scoring, GATK v.4.2.0 was also used to assess the reproducibility of variant scoring results. For Strelka2, BAM files with duplicate reads marked were processed with Manta (Chen et al., 2016; https://github.com/Illumina/manta) to obtain a list of candidate indel sites. After Manta processing, Strelka2 v. 2.9.10 was configured using the default exome mode and candidate calling regions obtained from Manta. Default filtering parameters were used. For FreeBayes variant caller we applied the default set of settings and filtered the resulting variant set by quality (QUAL < 30) and other recommended parameters using GATK v. 4.1.4.1.

### Benchmarking of variant discovery tools

Benchmarking was performed using the hap.py tool, a reference implementation of the GA4GH recommendations for variant caller benchmarking (Krusche et al., 2019). RTGtools vcfeval was used as an engine for comparison (Cleary et al., 2015). For all samples and variant discovery pipelines, performance was evaluated using a set of GENCODE v19 exon regions with an additional 150 bp padding upstream and downstream of each exon. For WES samples, an additional BED file was provided to limit the analysis to targeted exon regions that are included into the design as indicated by the kit vendor. A set of high-confidence variant calling regions was used to make all comparisons.

Reference stratification BED files were used for benchmarking alongside custom BED. Several custom sets of regions were added to this set: (i) regions upstream and downstream of each CDS sequence (0-25 bp, 25-50 bp, 50-75 bp, 75-100 bp, 100-125 bp, and 125-150 bp), regions with varying fraction of reads with MQ = 0 (multimapper fraction, Barbitoff et al., 2020), and regions with different expected normalized coverage obtained using a coverage model (Barbitoff et al., 2020).

### Statistical analysis

Statistical analysis of coverage statistics and benchmarking results was performed using R v. 4.2 with the following external packages: cowplot, colorRamps, ggplot2 (Wickham, 2016), ggsci (https://github.com/nanxstats/ggsci), lattice, reshape2. Statistical comparison between short read alignment software and variant callers were performed using the paired Wilcoxon signed rank test.

### Data availability

All data and code pertinent to the analysis presented here is available through GitHub:https://github.com/bioinf/caller_benchmark.

## Results

### Data collection and analysis strategy

To dissect the factors that define the accuracy of variant calling, we selected a matching set of WES and WGS datasets available for the gold standard GIAB samples, including NA12878 (HG001), three members of an Ashkenazi trio (HG002 - HG004), and one member of the Chinese Han trio (HG005). All WES datasets used in the analysis were prepared using the same capture kit (Agilent SureSelect All Exon v5), and all WGS samples corresponded to PCR-free WGS technology. The following analysis strategy was employed for each sample (Figure 1a): raw reads were aligned onto the GRCh37 human reference genome with either of the 4 short read aligners: Bowtie2, BWA MEM, Isaac, and Novoalign. Resulting alignment files in BAM format were subject to pre-processing with GATK to mark duplicate reads, and were then processed with 4 different variant calling software tools: DeepVariant, Strelka2, GATK-HC, and FreeBayes (see Methods for details). Raw variant calls were subject to filtering with standard built-in filters or quality-based filtering (see Methods), and filtered variant call sets were evaluated using the hap.py toolkit, with an additional stratification of coding regions by expected read depth, GC content, mappability, and other factors (see below).

**Figure 1.**
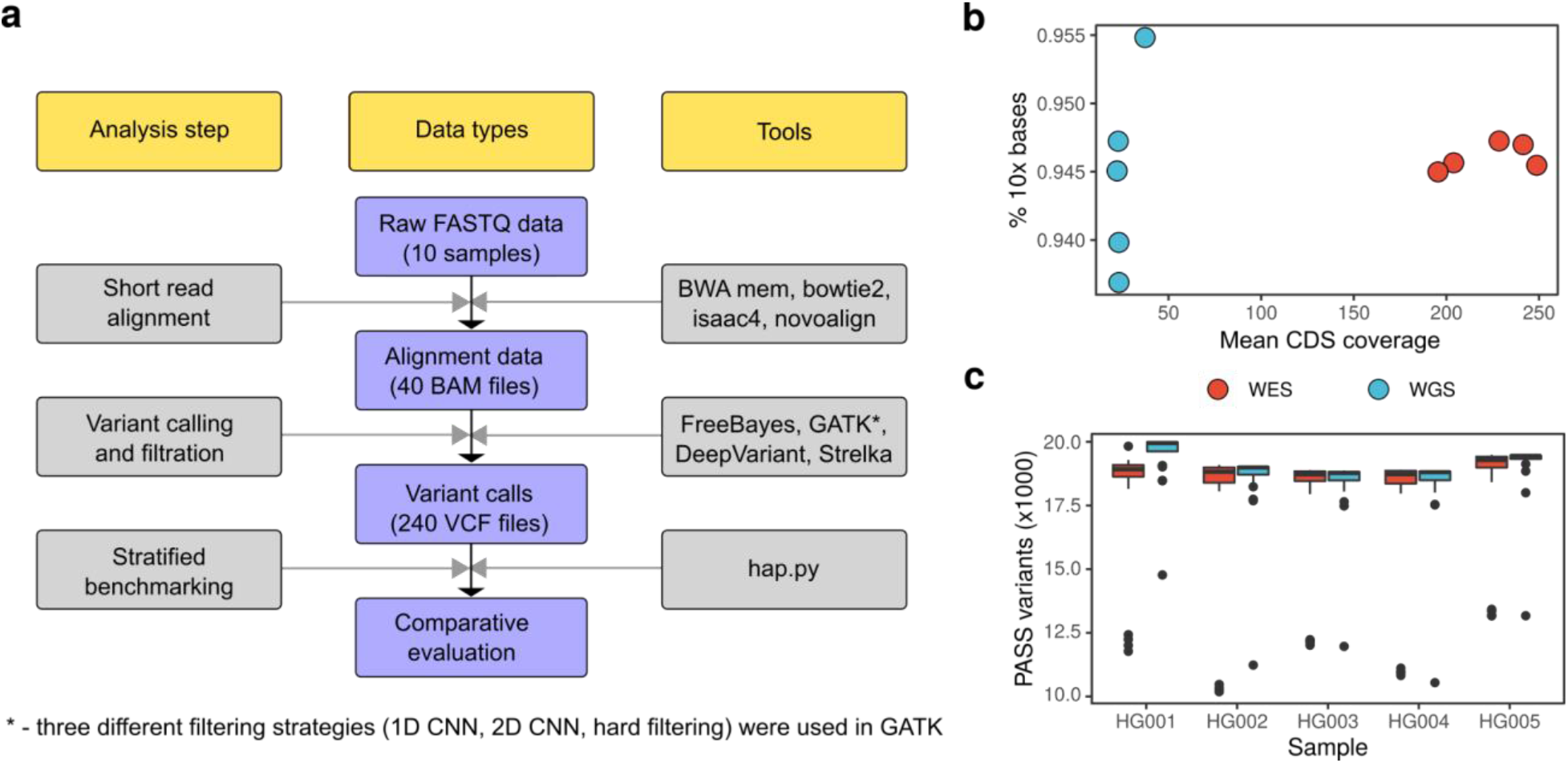
Systematic benchmarking of multiple variant calling pipelines. (a) A chart representing the analysis workflow. (b) A scatterplot showing mean coverage of high-confidence coding sequence regions (defined by the Genome In A Bottle consortium) and the fraction of bases of such regions covered at least 10x total read depth in WGS and WES datasets used (each point corresponds to an individual sample). (c) Number of pass-filter variants in each WES and WGS dataset discovered with all variant calling pipelines used in the study. Outliers correspond to pipelines which yield a substantial fraction of filtered variants.

Prior to the analysis of benchmarking results, we compared the overall quality and coverage of coding sequences in WES and WGS samples used. As the WES dataset for HG001 contained more than 250 million reads (Supplementary Figure 1a), we randomly selected 40% of all read pairs prior to analysis. All WES samples were characterized with significantly higher mean coverage of CDS regions and had comparable percentages of regions covered with at least 10 reads (Figure 1b, Table 1, Supplementary Figure 1c). At the same time, the fraction of bases with at least 20x coverage was higher in WES than in WGS (Supplementary Figure 1b). Overall, these results suggest that the difference in the estimated variant calling performance on WES and WGS data should not be driven by general low coverage in WES or by CDS regions not included into the WES capture kit design.

**Table 1.** Descriptive statistics of the gold standard sequencing datasets used in the study.

| Sample | Type | Source (SRA ID) | Mean coverage | Fraction of 10x bases | Median variant count** | True variant count |
| --- | --- | --- | --- | --- | --- | --- |
| HG001 | WGS | GIAB FTP | 22.2 | 0.937 | 19988 | 19997 |
|  | WES | ERR1905890 | 248.8* | 0.945* | 18971 |  |
| HG002 | WGS | GIAB FTP | 23.2 | 0.947 | 19039 | 19052 |
|  | WES | SRR2962669 | 241.4 | 0.945 | 18872 |  |
| HG003 | WGS | GIAB FTP | 23.2 | 0.948 | 18827 | 18920 |
|  | WES | SRR2962692 | 203.9 | 0.947 | 18770 |  |
| HG004 | WGS | GIAB FTP | 22.8 | 0.944 | 18857 | 18933 |
|  | WES | SRR2962694 | 228.4 | 0.940 | 18780 |  |
| HG005 | WGS | GIAB FTP | 37.3 | 0.955 | 19493 | 19501 |
|  | WES | SRR2962693 | 195.5 | 0.946 | 19333 |  |

Coverage and variant statistics are given with respect to GIAB high-confidence CDS regions. \* - coverage values for downsampled HG001 WES dataset are given (see Methods); \*\* - variant counts were obtained by calculating the median number of variants discovered by each pipeline.

We then evaluated the total number of variants discovered inside the high-confidence protein-coding regions with each of the 24 variant discovery pipelines used. All samples had a comparable number of variant calls, with a median value slightly above or below 19,000 variants per sample. In all cases, slightly fewer variants were discovered for WES samples compared to WGS (Table 1). For most samples the difference in variant count between WES and WGS was below 200 variants; however, for HG001 it was much more pronounced (1017 variants), most probably due to the differences in high-confidence region definition (see Methods). Surprisingly, we also found that some of the variant calling pipelines yield a very low number of pass-filter variants for both WES and WGS data (Figure 1c, outlier points). The reasons for such behavior will be discussed in detail later.

### Systematic comparison of short read alignment and variant calling software

Usage of a set of 10 independent sequencing datasets from 5 individuals allows us not only to compare the performance of different tools, but also to assess the robustness and reproducibility of variant caller performance. To conduct such an analysis, we first examined the F1 scores (a harmonic mean of precision and recall) using variant calls inside CDS regions generated by each combination of read alignment and variant calling software (Figure 2a). This analysis showed that variant callers seem to have a greater influence on the overall performance of a pipeline compared to short read aligners. Among all pipelines tested, a combination of BWA MEM with DeepVariant had the greatest F1 score, while DeepVariant showed best performance on both SNP and indels for any aligner. Among other solutions, Strelka2 showed high accuracy when working with BWA, Isaac, or Novoalign; at the same time, Strelka2 performance dramatically dropped when using Bowtie2 as the read aligner (Figure 2a). FreeBayes performed considerably worse than the aforementioned tools on both SNPs and indels, while GATK-HC had high accuracy only when 1D CNN or hard filtering strategy was used. GATK-HC combined with the 2D CNN variant filtering showed the worst performance in SNP calling irrespective of the aligner used. The reasons for this will be discussed in detail later.

**Figure 2.**
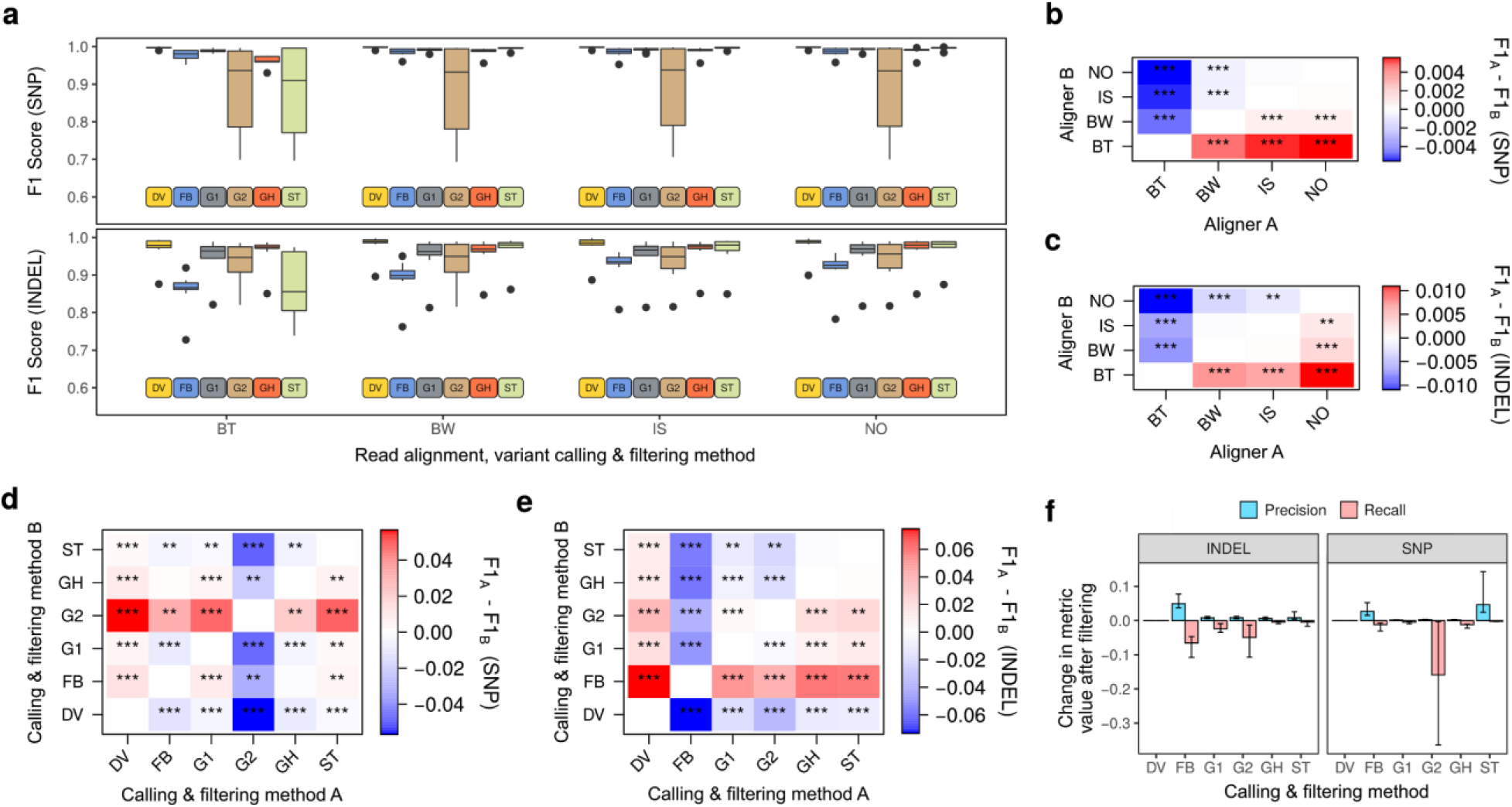
Statistical comparison of variant discovery pipelines’ performance. (A) Box plots representing the F1 scores for different combinations of aligners and variant callers. (B - E) Pairwise comparison of tool performance for read aligners (B, C) and variant callers (D, E) using pass-filter SNP (B, D) and indel (C, E) calls. (F) Difference in precision and recall values upon variant filtration with recommended settings. Read aligners: BW - BWA MEM, BT - Bowtie2, IS-isaac4, NO - Novoalign; variant callers and filtering strategies: DV - DeepVariant, G1 - GATK HaplotypeCaller with 1D CNN filtering, G2 - GATK HaplotypeCaller with 2D CNN filtering, GH-GATK HaplotypeCaller with recommended hard filters. ST - Strelka2, FB - FreeBayes.

We then sought to make a formal statistical comparison of read aligners and variant callers based on the benchmarking results. The structure of our dataset allows us to make such a comparison in a paired manner, and such formal paired-sample comparison could in turn provide important information on the reproducibility of the differences between pipelines on various datasets. Pairwise comparison of all short read aligners showed that, despite low median differences in F1 scores (maximum value of F1 difference for a pair of aligners was 0.0038 for SNP and 0.0104 for indels), Isaac and Novoalign show the best performance on both SNPs (Figure 2b) and indels (Figure 2c); they are closely followed by BWA MEM. Bowtie2, on the other hand, performed considerably worse with both SNPs and indel variants (Figure 2c). Among all variant calling and filtering solutions, DeepVariant showed better performance compared to all other tools (p-value < 0.001, Figure 2d, Figure 2e). Consistent with earlier observations, GATK-HC with 2D CNN model performed reproducibly worse on SNPs than any other pipeline. At the same time, FreeBayes was the worst solution for indel discovery, with its F1 score being 5.7% lower compared to any other method (Figure 2e). GATK-HC with hard filtering and Strelka2 performed almost equally well on SNPs and indels (p-value > 0.001 and > 0.05, respectively) and were the closest runners-up to DeepVariant. Taken together, our statistical comparison demonstrated that the performance differences for variant callers and read aligners are reproducible when using different input data. The results also confirm that read alignment differences have a generally lower impact on the accuracy of variant discovery compared to variant calling software.

We next questioned if the observed differences between variant calling pipelines can be mostly attributed to differences in precision or recall. To address this question, we compared precision and recall values reported for the same set of pipelines. This analysis revealed that almost all read aligners and variant callers, including the least efficient ones, had comparable precision values (with only a slight preference for DeepVariant for both SNPs and indels (p < 0.001) and a surprising benefit in precision from otherwise underperforming Bowtie2 (p < 0.01, Supplementary Figure S2). At the same time, the difference in recall was much more substantial, with pipelines showing the lowest F1 scores showing the worst recall values (Supplementary Figure S3). Given these observations, we conclude that the reproducible differences between variant calling software arises mostly from the power differences and not from different false positive rates.

Dramatic differences in recall values between variant calling pipelines prompted us to ask to what extent variant filtering negatively influences the results. To test this, we first compared the F1 scores for the raw unfiltered data (Supplementary Figure S4). Remarkably, we found that the differences between F1 scores of different pipelines on unfiltered data were much less dramatic, though tools that showed best or worst performance on filtered data tended to have higher or lower F1 values on unfiltered data as well. This result indicates that variant filtering might decrease recall more than increase precision. To formally test this hypothesis, we compared the F1 scores for variant sets before and after filtering with various filtering strategies. This analysis showed that, indeed, standard variant filtering for FreeBayes, as well as all filtering methods for GATK-HC, results in a higher median loss of recall compared to the median gain in precision (Figure 2f, Table 2). In contrast, standard filtering of variants in Strelka2 provided a reasonable gain in precision and a relatively low loss of recall. For DeepVariant, accurate comparison could not be made due to the lack of information about filtered genotypes.

**Table 2.** Effects of standard variant filtering methods on precision and recall.

| Caller and filtering strategy | Filtering effects on SNPs* |  |  | Filtering effects on indels* |  |  |
| --- | --- | --- | --- | --- | --- | --- |
|  | Raw calls F1 | Precision gain | Recall loss | Raw calls F1 | Precision gain | Recall loss |
| DeepVariant (default filter) | 0.998 | n.a,** | n.a.** | 0.989 | n.a.** | n.a.** |
| Strelka2 (default filter) | 0.955 | **0.046** | -0.002 | 0.972 | **0.009** | -0.004 |
| GATK (1D CNN, tranches 99.9/99.5) | 0.995 | 0.001 | **-0.005** | 0.975 | 0.008 | **-0.024** |
| GATK (2D CNN, tranches 99.9/99.5) |  | 0.003 | **-0.159** |  | 0.008 | **-0.049** |
| GATK (recommended hard filtering) |  | 0.002 | **-0.012** |  | **0.006** | -0.004 |
| Freebayes (standard quality-based filter) | 0.981 | **0.026** | -0.012 | 0.924 | 0.049 | **-0.065** |

\* - median values across all samples are shown, greater absolute value (precision gain/recall loss) for each filtering strategy is highlighted in bold; \*\* - effects of filtering on DeepVariant calls could not be assessed due to the structure of the output files.

To sum up, we demonstrate that variant calling pipelines reproducibly differ in their performance of variant discovery (both for SNPs and indels). These differences are mostly driven by sensitivity of variant caller software, which is greatly affected by variant filtration for most of the tested tools. Pipelines based on DeepVariant consistently perform better than all other solutions, while usage of Bowtie2 and FreeBayes is not recommended due to large losses in accuracy, especially in certain combinations.

### Analysis of factors influencing the accuracy of variant discovery

We next went on to evaluate the influence of sequencing technology and other sequencebased factors on the accuracy of variant discovery with different tools. To do so, we conducted a stratified benchmarking of variant discovery pipelines using hap.py with both pre-defined stratifications provided by GIAB and custom region sets obtained from the coverage model (Barbitoff et al., 2020).

We first compared the performance of different variant calling pipelines on WES and WGS data. Analysis of F1 score for SNPs and indels revealed that, while accuracy of SNP discovery was comparable for WES and WGS, indel variants are harder to call using exome data (Figure 1a, Supplementary Figure S5, Supplementary Figure S6). This result was reproducible across different read alignment and variant calling software, with the exception of FreeBayes which showed comparably low efficiency of indel calling with both WES and WGS data as input. Interestingly,

GATK-HC with the 2D CNN variant scoring performed much worse on WES than on WGS data; in particular, due to very low accuracy of SNP filtration (we were unable to fix such behavior with parameter tuning). At the same time, scoring variants with the 2D CNN provided substantially higher accuracy with WGS data. Hence, it can be concluded that the 2D scoring model should be applied only to whole-genome datasets with a more even coverage profile. Importantly, the aggregate differences between WES and WGS in SNP calling F1 scores for best variant calling pipelines were smaller compared to the differences in variant caller performance (median performance difference = 0.0003 for SNP and 0.016 for indels), suggesting that WES allows for a reasonably accurate variant discovery within CDS regions with the best performing variant calling solutions.

Given our previous work on WES/WGS comparison (Barbitoff et al., 2020), we hypothesized that lower WES performance in indel discovery was driven by regions in the vicinity of the CDS borders. To test this, we compared the median F1 score of variant discovery with different variant callers in regions located 25, 50, 75, 100, 125, or 150 bp away from the exonintron boundary. This comparison revealed that the performance of best variant calling pipelines (Novoalign + DeepVariant/Strelka2/GATK-HC-1D) on SNPs declined modestly with increasing the distance from CDS for all pipelines (Figure 3b, Supplementary Figure S6), and the F1 metric value was comparable for CDS boundary and regions located up to 50 bp upstream and downstream of each CDS region. At the same time, reliability of indel variant discovery rapidly decreased, and the overall accuracy of indel calling dropped significantly even at the distance of 25 bp from the CDS, and was lower than exome-wide F1 even at the exon boundary itself (Figure 3b, Supplementary Figure S6).

**Figure 3.**
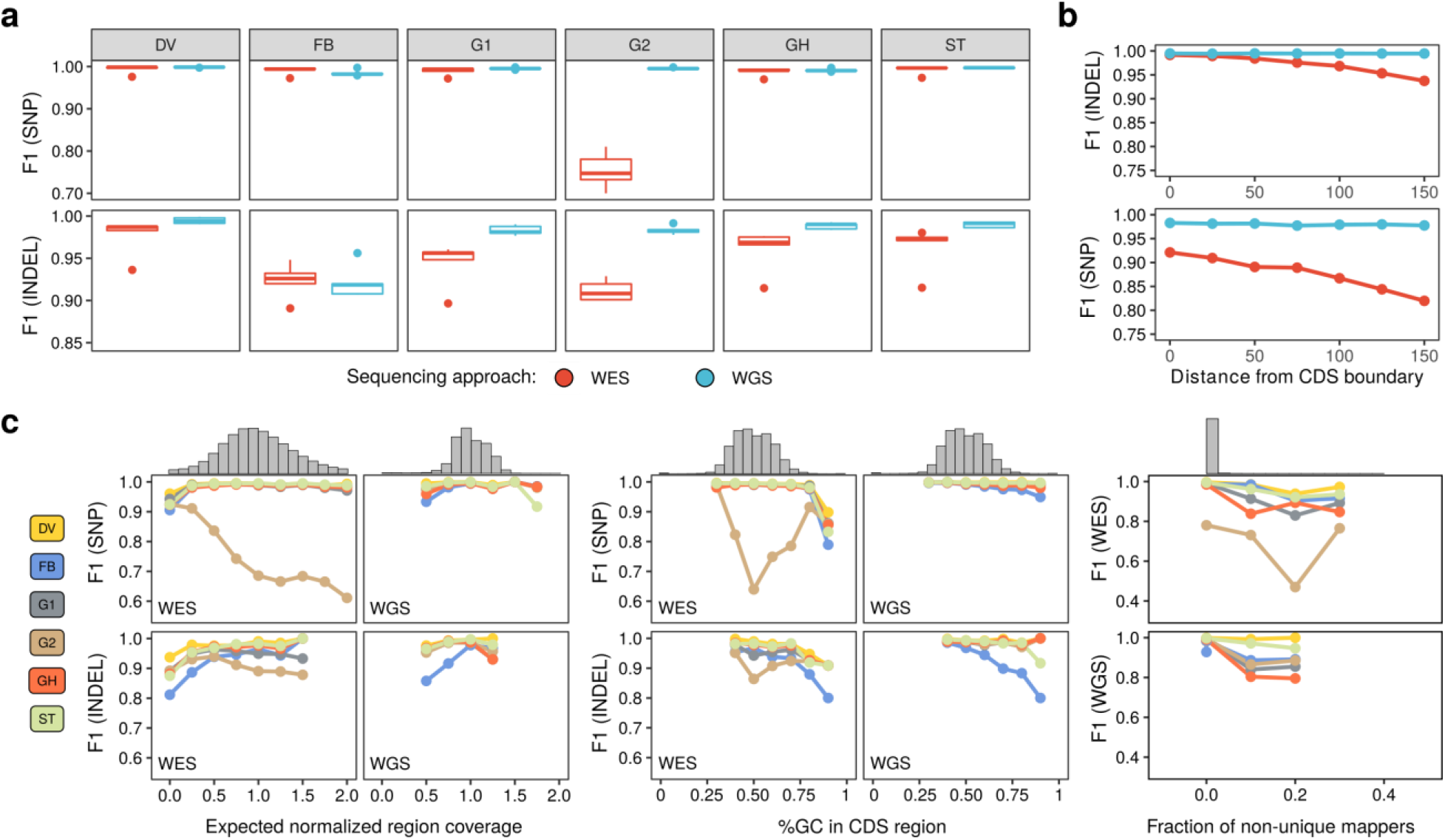
Dissection of factors affecting variant discovery pipelines’ performance. **(a)**Boxplots representing the F1 scores for SNP and indel calling with different variant callers in WES and WGS datasets. **(b)**The relationship between the distance from CDS boundary and the median F1 score of variant calling in WES and WGS data for SNP (top) and indel (bottom) variants. Results shown were obtained using the best-performing aligner (Novoalign) and three best-performing variant calling methods (DeepVariant, Strelka2, GATK-HC with hard filtering). **(c)**Comparison of the variant performance of variant callers in regions with different levels of expected normalized coverage (left), GC-content (middle), and fraction of non-unique mappers (right). Histograms on top of each plot represent the distribution of each parameter (coverage, GC content, MF) across GRCh37 CDS regions. Results shown on (a-c) were obtained using the best-performing read aligner (Novoalign v. 4.02.01). For other aligners, please see Supplementary Figures S5-S7. Variant callers and filtering strategies: DV - DeepVariant, G1 - GATK HaplotypeCaller with 1D CNN filtering, G2 - GATK HaplotypeCaller with 2D CNN filtering, GH - GATK HaplotypeCaller with recommended hard filters. ST - Strelka2, FB - FreeBayes.

Remarkably, variant calling pipelines that performed worse in the general comparison (Figure 2) also showed greater rate of the accuracy decay with increasing the distance from CDS boundary (Supplementary Figure S6). We were surprised to discover that the trend was reversed for the GATK-HC with 2D CNN filtering, which showed higher accuracy in regions more distant from the CDS boundary. High coverage might be one of the factors that negatively impacts the performance of this method. Taken together, our data suggest that variant calling in the regions flanking coding sequences (especially regions located no further than 50 bp away) is generally reliable for SNPs, but not indel variants when using WES data. Moreover, our observations suggest that the differences in indel calling accuracy between WES and WGS may be explained by low performance of variant callers near the exon boundaries. This offers an exciting opportunity to amend the probe design greatly improving this specific aspect of WES performance, bringing it even closer to the WGS.

We next turned to the analysis of other factors that determine the reliability of variant calling. There are plenty of sources of coverage bias in both WES and WGS experiments, as detailed previously (Barbitoff et al., 2020). However, it is unclear which sources of reproducible coverage bias in WES and WGS impact the performance of variant calling software. To assess this, we compared the performance of variant discovery in regions with systematic differences in normalized coverage, GC content, and the fraction of non-uniquely mapped reads (multimapper fraction, MF) (Barbitoff et al., 2020).

We started off by evaluating the dependence of the F1 score for each variant calling solution on the expected level of normalized sequence coverage as predicted by our recently proposed coverage model (Barbitoff et al., 2020). As expected, accuracy of all variant callers was decreased in regions with low normalized coverage in both WES and WGS. Surprisingly, we found that most variant calling pipelines also underperform in regions with high normalized coverage, especially for WES samples (Figure 3c, left panel; Supplementary Figure S7). Such a drop in performance in high-coverage regions is the most pronounced for GATK neural network-based filtering methods (the most dramatic effect is seen for the 2D CNN method, in line with earlier observations (Supplementary Figure S7)). Strelka2 is also slightly sensitive to high read depth, particularly with WGS datasets.

Yet another source of coverage biases in both WES and WGS is the GC-content of the sequence. We have previously shown that, while GC content is not a dominant determinant of poor sequencing coverage, extremely GC-rich or GC-poor regions tend to be covered substantially worse in WES. To assess the effects of the GC-content on the performance of variant calling software, we evaluated the F1 scores for each variant calling pipeline in regions with different GC content (split into 10% windows). We found that most variant callers’ performance drops significantly in extremely GC-rich regions (Figure 3c, middle; Supplementary Figure S7). Again, the effect of GC content on variant caller accuracy was the most pronounced for worst-performing variant calling methods such as FreeBayes; and the GC-content affects indel calling more than SNP calling in both WES and WGS. Importantly, variant calling in GC-rich regions was less efficient in WGS as well, though the relative drop in performance of variant callers in GC-rich regions is less significant for WGS than for WES (Figure 3c).

At the same time, extreme GC-rich and GC-poor regions span not more than 80 kb of human coding sequence. Previously we demonstrated that mappability limitations of short reads play a much more important role in poor sequencing coverage in both WES and WGS. The performance of variant callers in completely repetitive coding regions (for example, in duplicated genes) cannot be evaluated as these regions are mostly unreachable for short reads. On the other hand, there are multiple CDS regions with imperfect repeats that are only partially covered by multimapping reads. To assess whether variant caller performance in such regions is decreased, we compared the F1 scores for all variant calling pipelines in regions with different proportions of non-uniquely mapped reads. This analysis showed that, indeed, accuracy of variant discovery is compromised in such regions (Figure 3c, right panel). Similarly to previous comparisons, best-performing solutions such as DeepVariant or Strelka2 tend to be less sensitive to read mappability issues, while GATK is the most sensitive to read mapping ambiguity.

Taken together, our results demonstrate that, while coverage, GC-content, and mapping quality all affect accuracy of variant discovery in coding sequences, the best-performing variant calling pipelines are less sensitive to such confounding factors and perform better in all coding regions. At the same time, differences in the quality of variant discovery between WES and WGS are subtle and are mostly attributable to the low power of indel calling near the exon boundaries.

## Discussion

NGS methods have dramatically transformed the world of human genetics, both from the research and clinical perspective. WES and WGS are becoming the new standard in the diagnosis of Mendelian disease. However, despite the rapid spread and wide application of NGS-based methods in clinical practice, the average diagnostic rate is still below 50% for both trio-based WGS and trio-based WES (Wright et al., 2018). Such low diagnostic rates are explained by a multitude of confounding factors that include inherent limitations of sequencing technologies, imperfect human reference genome sequence, software limitations and, perhaps most importantly, incomplete understanding of the disease etiology and pathology (Biesecker et al., 2019). Some of the inherent problems of the human reference genome, as well as limitations of WES and WGS, have been discussed elsewhere (Barbitoff et al., 2018; Ballouz et al., 2019; Ebbert et al., 2019; Barbitoff et al., 2020). In this study, we addressed the other major factor that plays an important role in variant discovery, namely, the performance of software pipelines for variant calling.

In contrast to other previously published studies, we undertook a more systematic approach by evaluating the performance of all different combinations of read aligners and variant callers using a large set of 5 WGS and WES samples. Such an approach allowed us to make estimates of the relative importance of different factors for accurate and reliable variant discovery. We show that variant calling software is the most important factor that greatly affects both SNP and indel calling. At the same time, the sequencing method (WGS or WES) has significant influence on the accuracy of indel detection, but virtually does not affect SNP calling (Table 3). Finally, read alignment software has generally the lowest influence on the accuracy of variant discovery in coding sequences, although the usage of Bowtie2 is discouraged (Figure 2b-c). Taken together, these results highlight the fact that the correct choice of software tools for variant discovery is paramount for high-quality variant calling.

**Table 3.** Aggregated median statistics of variant caller performance on WES and WGS data

| Caller (filtering)* | Type | SNP F1 | SNP Precision | SNP Recall | indel F1 | indel Precision | indel Recall |
| --- | --- | --- | --- | --- | --- | --- | --- |
| DeepVariant | WGS | **0.998587** | 0.998241 | **0.998934** | **0.992138** | **0.993377** | 0.991886 |
| Strelka2 | WGS | 0.997224 | 0.997916 | 0.996663 | 0.989820 | 0.982022 | **0.993304** |
| GATK (1D) | WGS | 0.995323 | 0.996901 | 0.994350 | 0.980811 | 0.988713 | 0.978541 |
| GATK (HF) | WGS | 0.990370 | 0.996921 | 0.984514 | 0.988789 | 0.991071 | 0.987830 |
| GATK (2D) | WGS | 0.995188 | 0.997321 | 0.993871 | 0.981703 | 0.990909 | 0.975391 |
| FreeBayes | WGS | 0.982014 | **0.998787** | 0.967933 | 0.925216 | 0.984655 | 0.875536 |
| DeepVariant | WES | **0.998270** | **0.997785** | **0.998756** | **0.986025** | **0.990762** | **0.976905** |
| Strelka2 | WES | 0.996510 | 0.996972 | 0.997133 | 0.972968 | 0.985782 | 0.962441 |
| GATK (1D) | WES | 0.993713 | 0.995222 | 0.991081 | 0.958550 | 0.973558 | 0.938967 |
| GATK (HF) | WES | 0.991012 | 0.997001 | 0.988495 | 0.969746 | 0.974359 | 0.967136 |
| GATK (2D) | WES | 0.747232 | 0.996936 | 0.597992 | 0.916065 | 0.974619 | 0.856808 |
| FreeBayes | WES | 0.994001 | 0.997716 | 0.992265 | 0.931814 | 0.969773 | 0.896714 |

All values are given with respect to the Novoalign v. 4.2.0 read alignment. Bold font corresponds to the best values for WGS and WES data. \* 1D - 1D CNN model in GATK, 2D - 2D CNN model in GATK, HF - hard filtering with recommended parameters.

Furthermore, we showed that variant callers mostly differ in their sensitivity (Supplementary Figures S2-S3) and the ability to accurately call variants in regions with poor sequencing coverage, extremely high or low GC-content, and/or non-zero fraction of multimappers (Figure 3; Supplementary Figure S7). We demonstrate that the most recently developed variant calling tools, such as DeepVariant (Poplin et al., 2018) and Strelka2 (Kim et al., 2018), have the highest robustness and provide high accuracy of variant discovery for all datasets used. These data support and expand previous observations that were made using individual gold standard samples (Supernat et al., 2018; Zhao et al., 2020). The best-performing solutions also tend to be less sensitive to these confounding factors such as depth of coverage and GC-content. Thus, our data suggest that recent developments in the field of variant calling software compensate for many of the limitations of short-read sequencing.

The DeepVariant method, the one that consistently shows the best accuracy of variant calling for both SNPs and indels, is based on a convolutional neural network model. Neural networks and other complex machine learning approaches are clearly the most promising for future development of bioinformatic software, including variant callers (Eraslan et al., 2019). At the same time, in some cases (e.g., GATK CNNScoreVariants tool) neural networks are more sensitive to artifacts and data quality, including sequencing depth (Figure 2, Figure 3). This problem likely arises from overfitting, *i.e.* excessive tuning of the model to show best performance on the specific sets of training data. Unfortunately, machine learning models for variant calling and filtering have been trained using the same GIAB gold standard sequencing datasets that are usually used for benchmarking (including in this study). This complicates the unbiased evaluation of the performance of these tools. It can be expected that deep learning-based callers and filtering methods would perform worse in non-GIAB quality datasets from human populations not included in GIAB. For this reason we would advocate the inclusion of a more diverse set of samples into the GIAB dataset, especially of African, Hispanic, or mixed ancestry. This would improve the models and increase robustness of the best-performing variant callers on different ethnical backgrounds, and increase the opportunity for cross-benchmarking. Recently, a new set of high-quality reference datasets have been generated for benchmarking of variant callers (Baid et al., 2020), and evaluation of several variant callers’ performance on these data corroborates the results present in our study. However, the new dataset is also based on the same individual GIAB genomes.

Given the results of our comparison, a simple set of recommendations could be made. Best-performing variant callers (DeepVariant, Strelka2, and GATK HC 1D) could all be used depending on the specifics (for example, DeepVariant was the slowest method used in our work, while Strelka2 was the fastest). At the same time, we show that variant filtering is not always done in an optimal way, and that filtered variants should be retained and carefully examined for medical genetic applications. It should also be noted that even the best variant callers could not make the correct call when the alignment is wrong. Thus, we would discourage the use of Bowtie2 which showed markedly lower performance in our benchmark. Finally, aside from a notable outlier of HG001, we confirm our previous conclusions that best WES solutions achieve performance that is very close to that of WGS, and should be considered a reliable and low cost option for many applications. Some of the calling and filtering methods, however, are incompatible with WES data and should only be applied to WGS.

While we believe that our results provide important insights into the performance of variant calling methods and highlight perspectives for future development, several important issues have not been explicitly addressed in our work and may require further exploration. First, the version of the human reference genome assembly also influences the accuracy of variant discovery, beyond the problem of reference minor alleles reported previously (Barbitoff et al., 2018). Apart from major differences between GRCh37 (used in this study) and GRCh38 reference assemblies, inclusion of unplaced contigs, patches, and decoy sequences could influence both quantity and quality of variant calls. Indeed, if compared side-by-side, different versions of the same human reference genome assembly yield different results of benchmarking with the same variant calling pipeline (Supplementary Table S2). Second, only high-confidence variant calling regions are routinely used for benchmarking of variant calling pipelines. While usage of such regions is useful to avoid the negative effects of coverage bias, it makes it harder to accurately compare variant caller sensitivity to low coverage, read mapping quality, and sequence complexity. Construction of a broader set of ground truth variant calls in non-high confidence regions would be useful for better benchmarking of variant calling methods, and for the development of robust new solutions for variant discovery in complex regions. Third, we did not address the differences that could be introduced by the specific short-read sequencing device. However, a recent publication comparing NovaSeq 6000, HiSeq 4000, MGISEQ-2000, and BGISEQ-500 (Chen et al., 2019) has found the differences to be modest, at least between the Illumina machines.

Finally, software tools for bioinformatic analysis of NGS data are constantly improving. Besides accuracy, running time (which was not specifically evaluated in our analysis) may also present a serious problem when working with large genomic datasets. Multiple attempts have been made recently to achieve high scalability of the read alignment and variant calling software. These include, but are not limited to, development of a native Google Cloud Platform integration in the recent versions of GATK, faster reimplementation of the BWA MEM algorithm (BWA-MEM2, Vasimuddin et al., 2019), and many others. New variant calling methods are also being developed, with Octopus (Cooket et al., 2021) being one of the most recent solutions. Constant development of novel methods and software tools suggests that large-scale stratified comparisons, like the one presented in our work, should be repeatedly conducted at least once in several years.

## Acknowledgements

This work was supported by the Systems Biology Fellowship to Y.A.B. the Presidential Fellowship for Young Scientists (grant no. SP-4503.2021.4) to Y.A.B., and D.O. Ott Research Institute of Obstetrics, Gynaecology and Reproductology, project 558-2019-0012 (AAAA-A19119021290033-1) of FSBSI. The authors declare no conflict of interest.

## References

Baid G, Nattestad M, Kolesnikov A, Goel S, Yang H, Chang PC, Carroll A. 2020. An extensive sequence dataset of gold-standard samples for benchmarking and development. bioRxiv. doi: 10.1101/2020.12.11.422022

Ballouz S, Dobin A, Gillis JA. 2019. Is it time to change the reference genome? Genome Biol 20:159.

Barbitoff YA, Bezdvornykh I V., Polev DE, Serebryakova EA, Glotov AS, Glotov OS, Predeus A V. 2018. Catching hidden variation: systematic correction of reference minor allele annotation in clinical variant calling. Genet Med 20:360–364.

Barbitoff YA, Polev DE, Glotov AS, Serebryakova EA, Shcherbakova I V., Kiselev AM, Kostareva AA, Glotov OS, Predeus A V. 2020. Systematic dissection of biases in whole-exome and whole-genome sequencing reveals major determinants of coding sequence coverage. Sci Rep 10:2057. doi: 10.1038/s41598-020-59026-y.

Biesecker LG, Green RC. 2014. Diagnostic Clinical Genome and Exome Sequencing. N Engl J Med 370:2418–2425.

Bycroft C, Freeman C, Petkova D, Band G, Elliott LT, Sharp K, Motyer A, Vukcevic D, Delaneau O, O’Connell J, Cortes A, Welsh S, et al. 2018. The UK Biobank resource with deep phenotyping and genomic data. Nature 562:203–209.

Chen X, Schulz-Trieglaff O, Shaw R, Barnes B, Schlesinger F, Källberg M, Cox AJ, Kruglyak S, Saunders CT. 2016. Manta: Rapid detection of structural variants and indels for germline and cancer sequencing applications. Bioinformatics 32:1220–1222.

Chen J, Li X, Zhong H, Meng Y, Du H. 2019. Systematic comparison of germline variant calling pipelines cross multiple next-generation sequencers. Sci Rep 9:1–13.

Cleary J, Braithwaite R, Gaastra K, Hilbush B, Inglis S, Irvine S, Jackson A, Littin R, Rathod M, Ware D, Zook J, Trigg L, et al. 2015. Comparing Variant Call Files for Performance Benchmarking of Next-Generation Sequencing Variant Calling Pipelines. bioRxiv 023754.

Cooke DP, Wedge DC, Lunter G. 2021. A unified haplotype-based method for accurate and comprehensive variant calling. Nat Biotechnol. In press doi: 10.1038/s41587-021-00861-3.

DePristo MA, Banks E, Poplin R, Garimella K V, Maguire JR, Hartl C, Philippakis AA, Angel G del, Rivas MA, Hanna M, McKenna A, Fennell TJ, et al. 2011. A framework for variation discovery and genotyping using next-generation DNA sequencing data. Nat Genet 43:491–498.

Ebbert MTW, Jensen TD, Jansen-West K, Sens JP, Reddy JS, Ridge PG, Kauwe JSK, Belzil V, Pregent L, Carrasquillo MM, Keene D, Larson E, et al. 2019. Systematic analysis of dark and camouflaged genes reveals disease-relevant genes hiding in plain sight. Genome Biol 20:97.

Eraslan G, Avsec Ž, Gagneur J, Theis FJ. 2019. Deep learning: new computational modelling techniques for genomics. Nat Rev Genet 20:389–403.

Garrison E, Marth G. 2012. Haplotype-based variant detection from short-read sequencing. arXiv:1207.3907

Hwang S, Kim E, Lee I, Marcotte EM. 2015. Systematic comparison of variant calling pipelines using gold standard personal exome variants. Sci Rep 5:17875. doi: 10.1038/srep17875.

Karczewski KJ, Francioli LC, Tiao G, Cummings BB, Alföldi J, Wang Q, Collins RL, Laricchia KM, Ganna A, Birnbaum DP, Gauthier LD, Brand H, et al. 2020. The mutational constraint spectrum quantified from variation in 141,456 humans. Nature 581:434–443.

Kim S, Scheffler K, Halpern AL, Bekritsky MA, Noh E, Källberg M, Chen X, Kim Y, Beyter D, Krusche P, Saunders CT. 2018. Strelka2: fast and accurate calling of germline and somatic variants. Nat Methods 15:591–594.

Krusche P, Trigg L, Boutros PC, Mason CE, La Vega FM De, Moore BL, Gonzalez-Porta M, Eberle MA, Tezak Z, Lababidi S, Truty R, Asimenos G, et al. 2019. Best practices for benchmarking germline small-variant calls in human genomes. Nat Biotechnol 37:555–560.

Langmead B, Salzberg SL. 2012. Fast gapped-read alignment with Bowtie 2. Nat Methods 9:357–359.

Li H, Durbin R. 2009. Fast and accurate short read alignment with Burrows-Wheeler transform. Bioinformatics 25:1754–1760.

Li H, Handsaker B, Wysoker A, Fennell T, Ruan J, Homer N, Marth G, Abecasis G, Durbin R. 2009. The Sequence Alignment/Map format and SAMtools. Bioinformatics 25:2078–2079.

McKenna A, Hanna M, Banks E, Sivachenko A, Cibulskis K, Kernytsky A, Garimella K, Altshuler D, Gabriel S, Daly M, Depristo MA. 2010. The Genome Analysis Toolkit: A MapReduce framework for analyzing next-generation DNA sequencing data. Genome Res 10:1297–1303.

Poplin R, Chang PC, Alexander D, Schwartz S, Colthurst T, Ku A, Newburger D, Dijamco J, Nguyen N, Afshar PT, Gross SS, Dorfman L, et al. 2018. A universal snp and small-indel variant caller using deep neural networks. Nat Biotechnol 36:983.

Supernat A, Vidarsson OV, Steen VM, Stokowy T. 2018. Comparison of three variant callers for human whole genome sequencing. Sci Rep 8:17851. doi: 10.1038/s41598-018-36177-7.

van der Auwera GA, Carneiro MO, Hartl C, Poplin R, Angel G del, Levy-Moonshine A, Jordan T, Shakir K, Roazen D, Thibault J, Banks E, Garimella K V., et al. 2013. From FastQ Data to High-Confidence Variant Calls: The Genome Analysis Toolkit Best Practices Pipeline. Curr Protoc Bioinforma 10.1–10.33.

van Dijk EL Auger H, Jaszczyszyn Y, Thermes C. 2014. Ten years of next-generation sequencing technology. Trends Genet 30:418–426.

Vasimuddin M, Misra S, Li H, Aluru S. 2019. Efficient Architecture-Aware Acceleration of BWA-MEM for Multicore Systems.

Wickham H. 2016. Ggplot2: Elegant Graphics for Data Analysis. 260 p.

Wright CF, FitzPatrick DR, Firth H V. 2018. Paediatric genomics: diagnosing rare disease in children. Nat Rev Genet 19:253–268.

Zook JM, Catoe D, McDaniel J, Vang L, Spies N, Sidow A, Weng Z, Liu Y, Mason CE, Alexander N, Henaff E, McIntyre ABR, et al. 2016. Extensive sequencing of seven human genomes to characterize benchmark reference materials. Sci Data 3:160025.

Zhao S, Agafonov O, Azab A, Stokowy T, Hovig E. 2020. Accuracy and efficiency of germline variant calling pipelines for human genome data. Sci Rep 10:20222. doi: 10.1038/s41598-020-77218-4

