## Supplementary Figures for "Systematic benchmark of state-of-the-art variant calling pipelines identifies major factors affecting accuracy of coding sequence variant discovery"

**Supporting information**

**Supplementary Files**

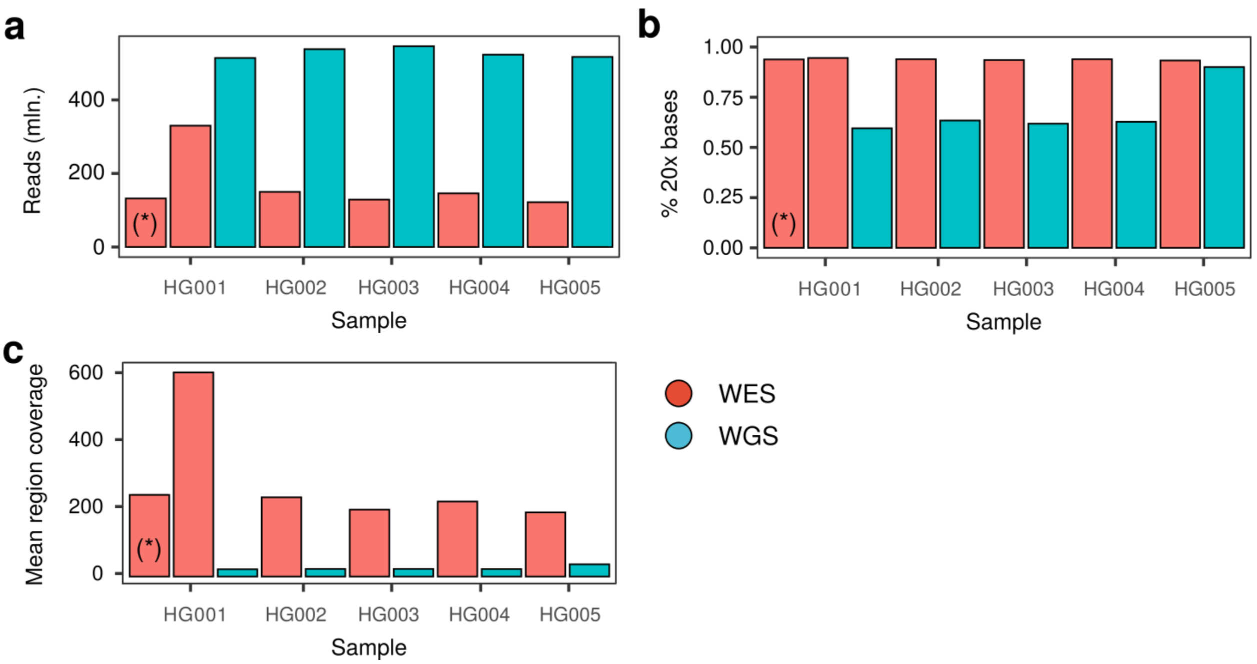


**Figure S1.** Major metrics of the datasets used in the benchmarking analysis. (a) Total number of reads per sample, (b) percentage of high-confidence CDS bases (defined by GIAB) covered at at least 20x depth. (c) mean coverage of high-confidence CDS regions (*) - downsampled dataset that was used throughout the analysis.

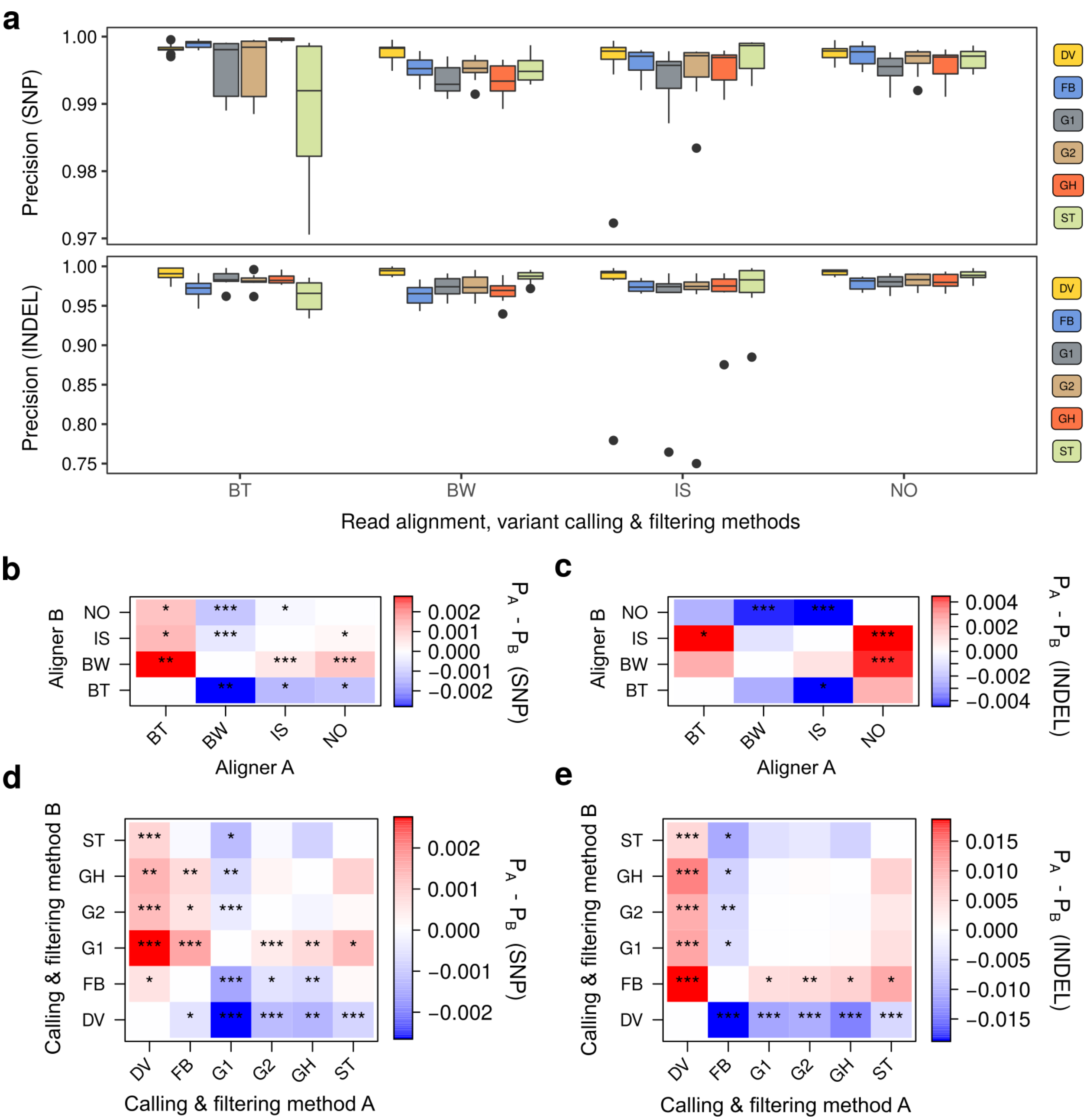


**Figure S2.** Comparison of the precision of alignment and variant calling methods. Top, box plots representing precision values for SNP and indel calling for all combinations of read alignment and variant calling software. Bottom, matched pairwise comparison of read alignment or variant callers in terms of precision of SNP and indel discovery. Read aligners: BW - BWA MEM, BT - Bowtie2, IS - isaac4, NO - Novoalign; variant callers and filtering strategies: DV - DeepVariant, G1 - GATK HaplotypeCaller with 1D CNN filtering, G2 - GATK HaplotypeCaller with 2D CNN filtering, GH - GATK HaplotypeCaller with recommended hard filters. ST - Strelka2, FB - Freebayes.

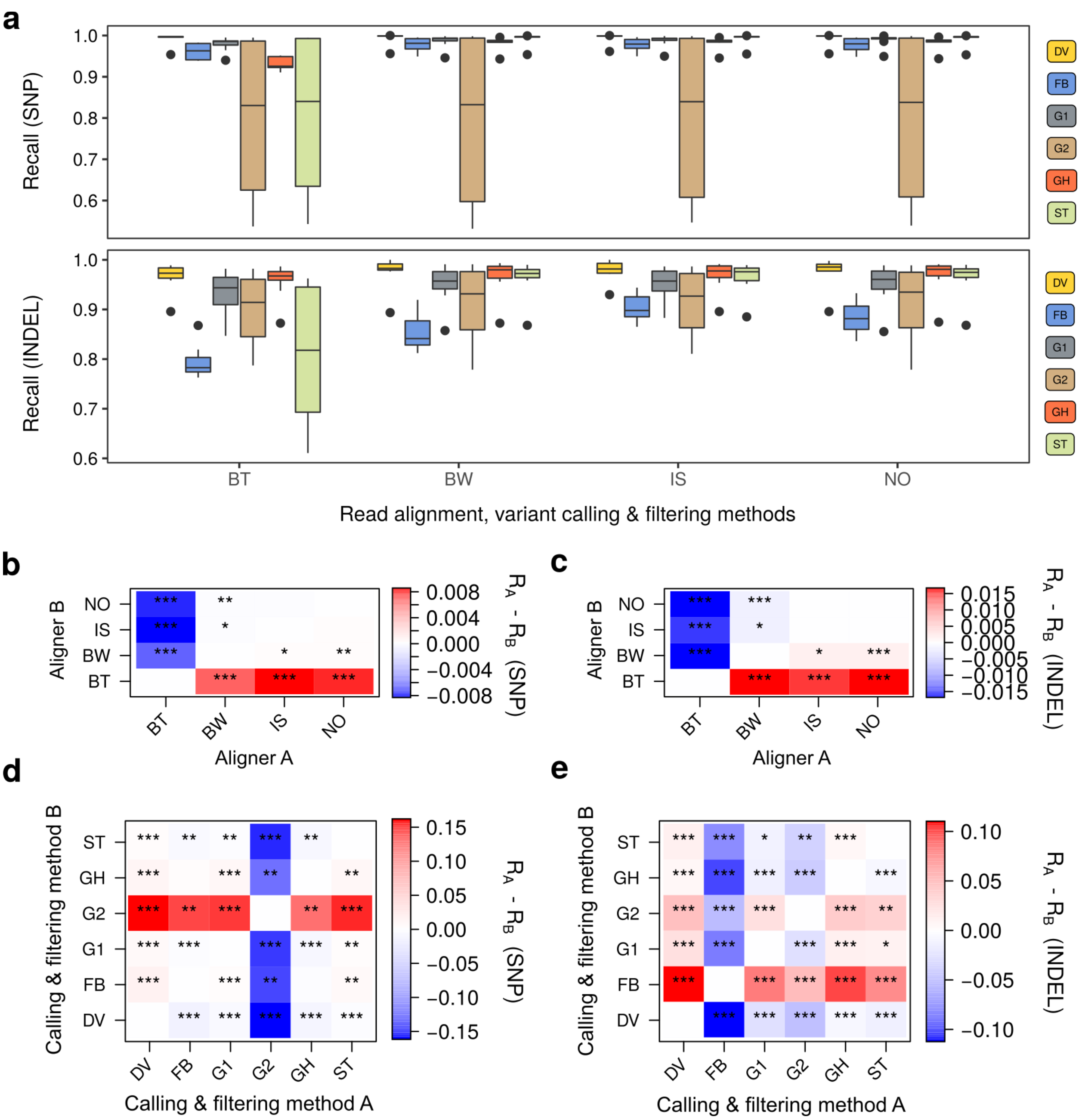


**Figure S3.** Comparison of the recall of alignment and variant calling methods. Top, box plots representing precision values for SNP and indel calling for all combinations of read alignment and variant calling software. Bottom, matched pairwise comparison of read alignment or variant callers in terms of precision of SNP and indel discovery. Read aligners: BW - BWA MEM, BT - Bowtie2, IS - isaac4, NO - Novoalign; variant callers and filtering strategies: DV - DeepVariant, G1 - GATK HaplotypeCaller with 1D CNN filtering, G2 - GATK HaplotypeCaller with 2D CNN filtering, GH - GATK HaplotypeCaller with recommended hard filters. ST - Strelka2, FB - Freebayes.

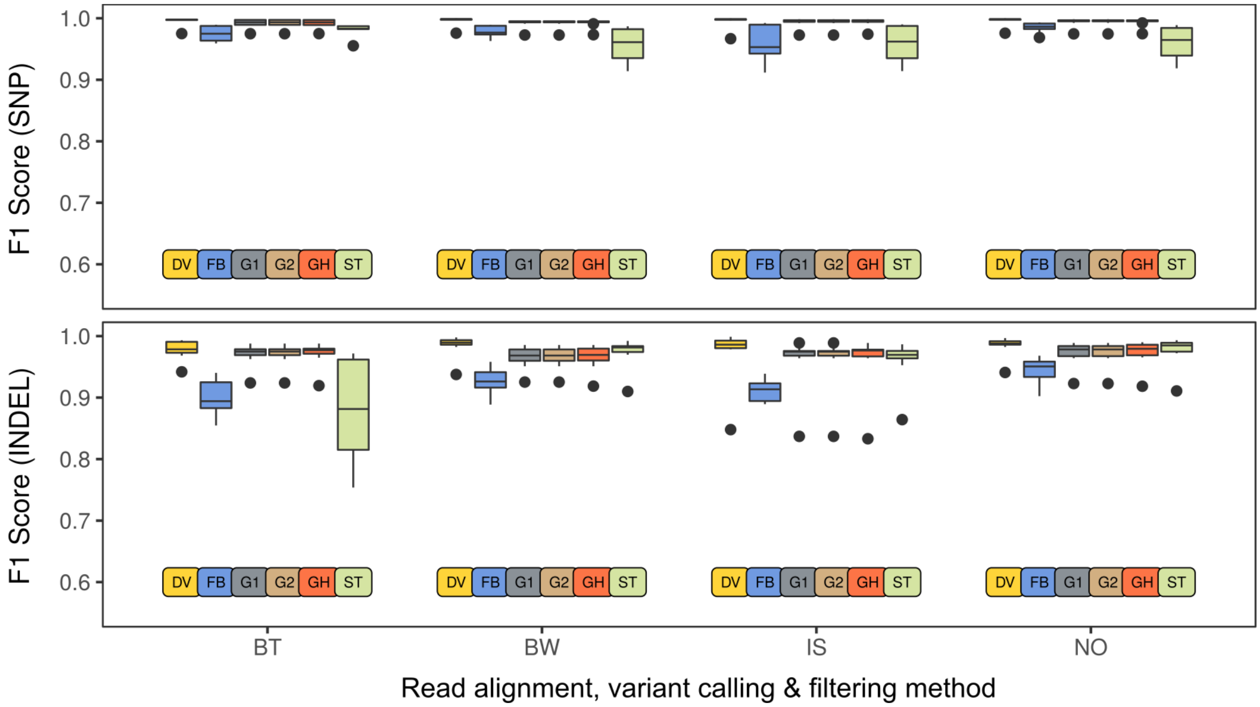


**Figure S4.** Comparison of the F1 scores of variant calling pipeline prior to applying filters. Please note that all pipelines based on GATK have identical F1 scores before filtering. Read aligners: BW - BWA MEM, BT - Bowtie2, IS - isaac4, NO - Novoalign; variant callers and filtering strategies: DV - DeepVariant, G1 - GATK HaplotypeCaller with 1D CNN filtering, G2 - GATK HaplotypeCaller with 2D CNN filtering, GH - GATK HaplotypeCaller with recommended hard filters. ST - Strelka2, FB - Freebayes.

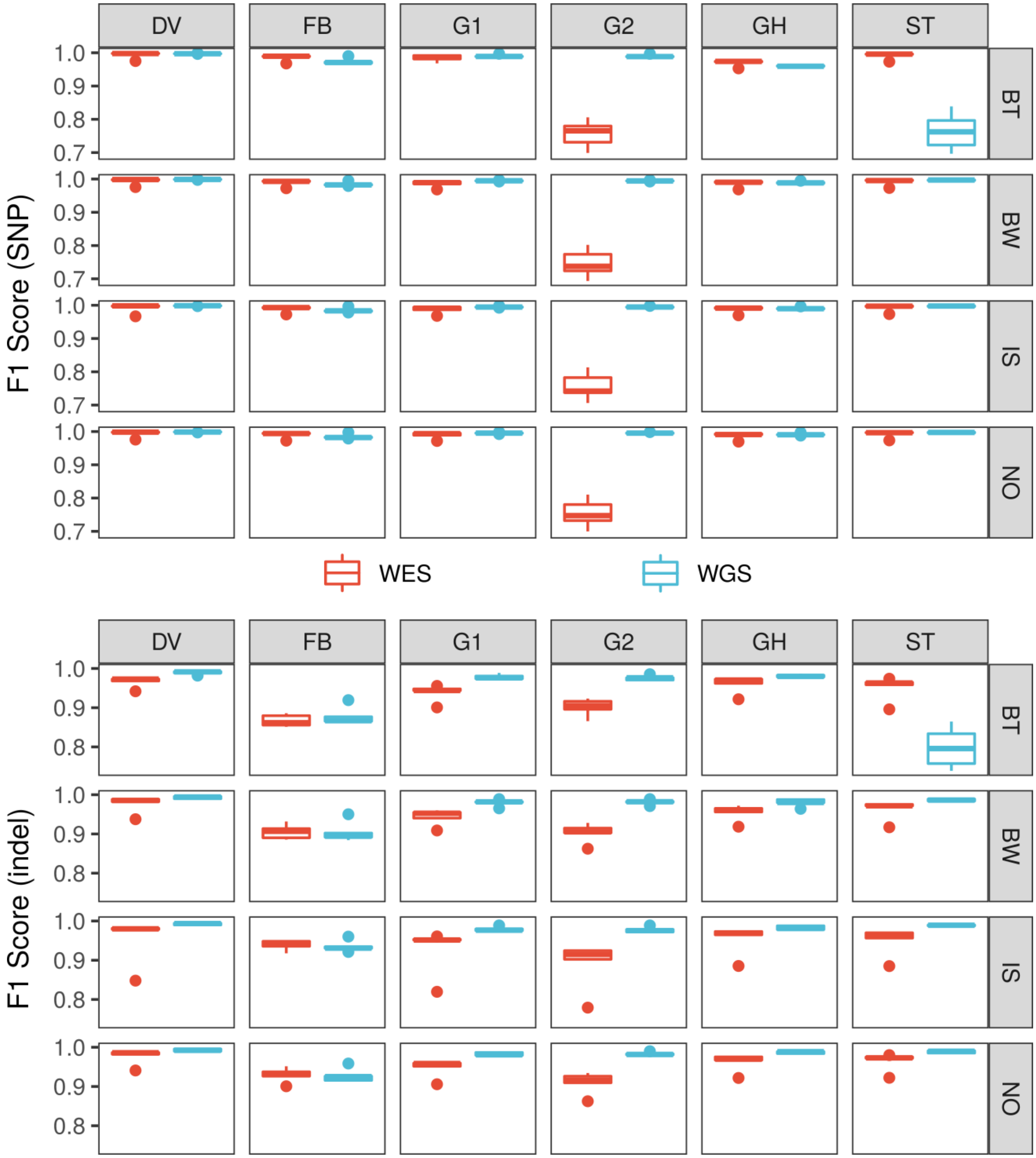


**Figure S5.** Comparison of variant calling pipelines' performance on WES and WGS data. F1 scores for SNP (top) or indel (bottom) calling are shown. Read aligners: BW - BWA MEM, BT - Bowtie2, IS - isaac4, NO - Novoalign; variant callers and filtering strategies: DV - DeepVariant, G1 - GATK HaplotypeCaller with 1D CNN filtering, G2 - GATK HaplotypeCaller with 2D CNN filtering, GH - GATK HaplotypeCaller with recommended hard filters. ST - Strelka2, FB - Freebayes.

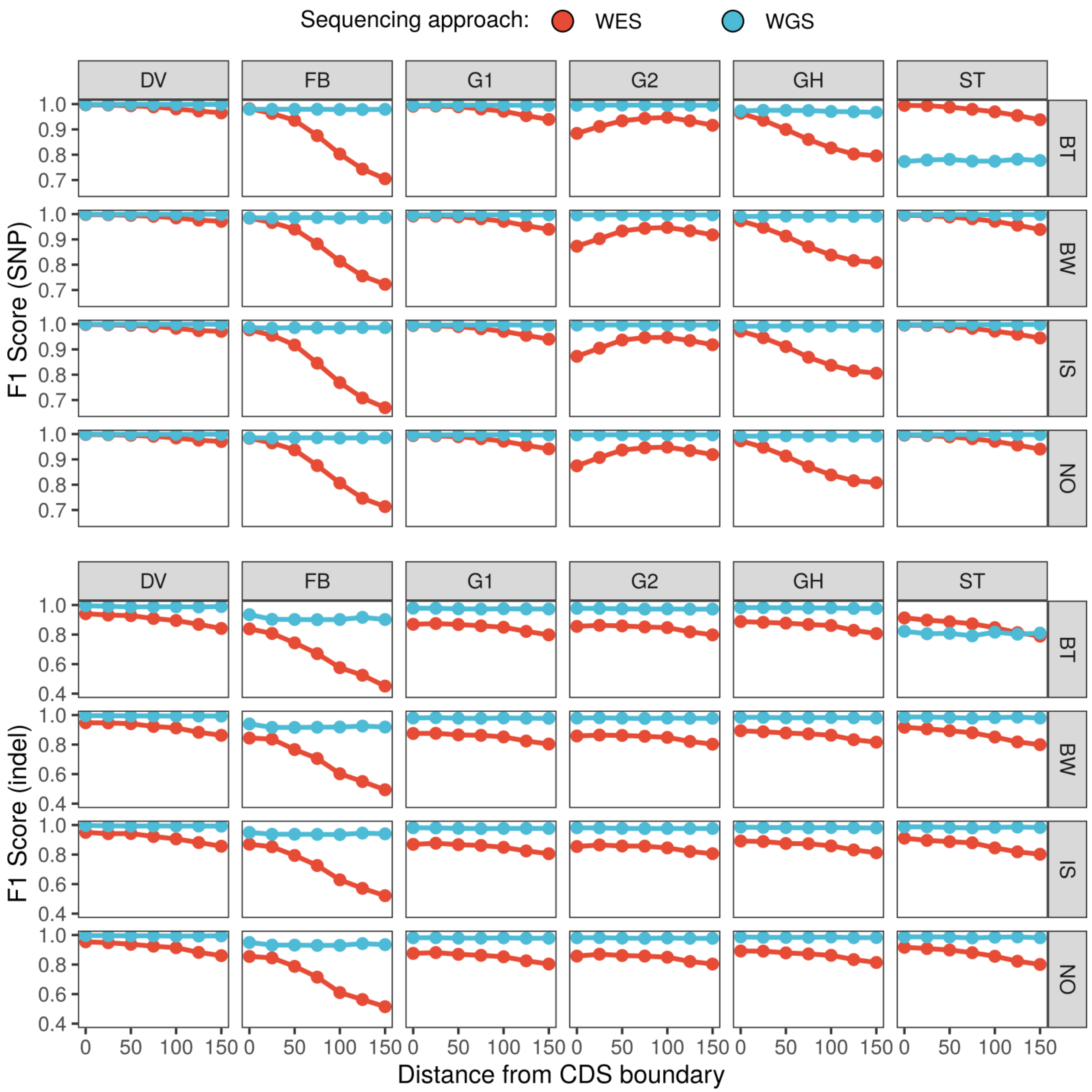


**Figure S6.** Accuracy of variant discovery in the vicinity of CDS regions for all variant calling pipelines. F1 scores for SNP (top) and indel (bottom) calling are shown. Read aligners: BW - BWA MEM, BT - Bowtie2, IS - isaac4, NO - Novoalign; variant callers and filtering strategies: DV - DeepVariant, G1 - GATK HaplotypeCaller with 1D CNN filtering, G2 - GATK HaplotypeCaller with 2D CNN filtering, GH - GATK HaplotypeCaller with recommended hard filters. ST - Strelka22, FB - Freebayes.

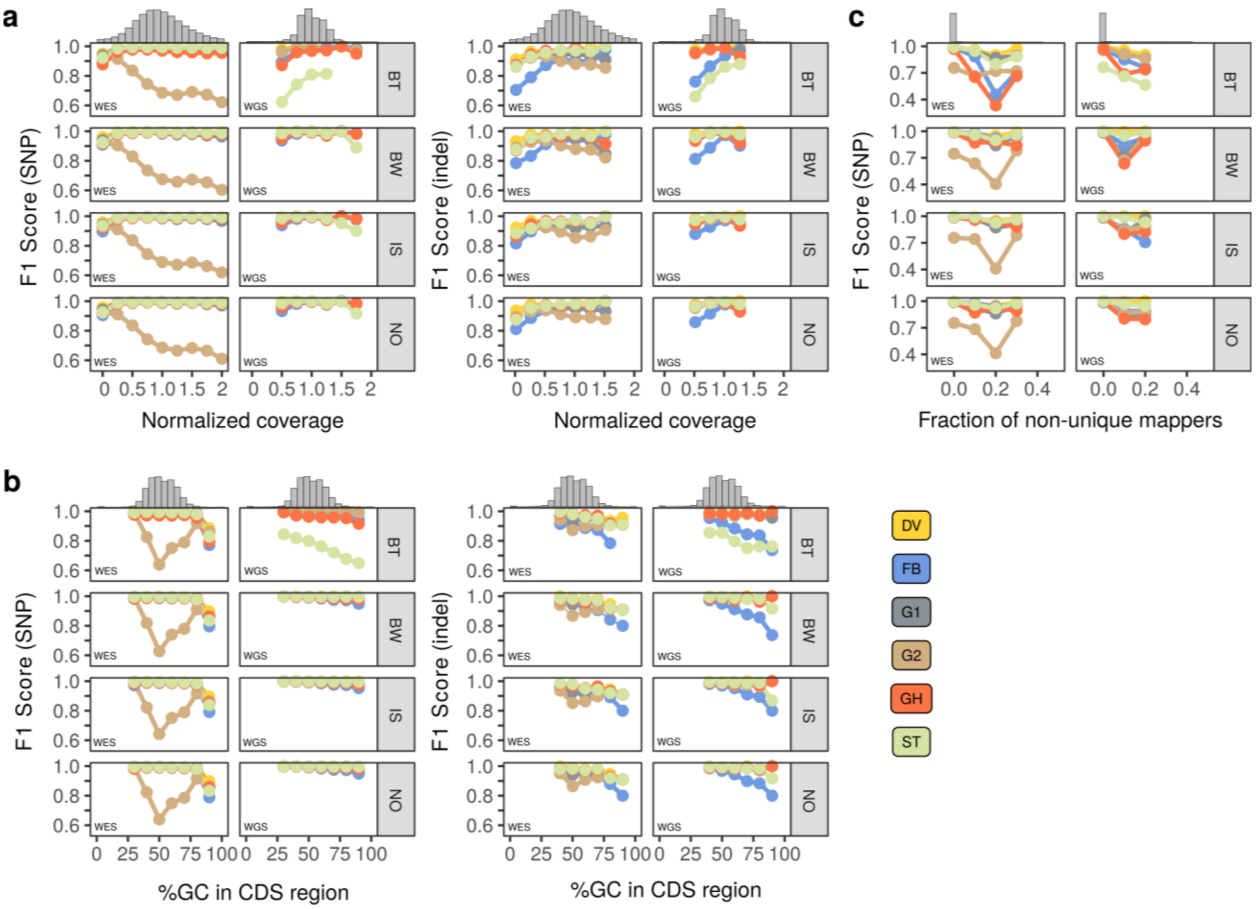


**Figure S7.** Comparison of the variant calling pipelines' performance in regions with different levels of expected normalized coverage (a), GC-content (b), and fraction of non-unique mappers (c). Histograms on top of each plot represent the distribution of each parameter (coverage, GC content, MF) across GRCh37 CDS regions. Read aligners: BW - BWA MEM, BT - Bowtie2, IS - isaac4, NO - Novoalign; variant callers and filtering strategies: DV - DeepVariant, G1 - GATK HaplotypeCaller with 1D CNN filtering, G2 - GATK HaplotypeCaller with 2D CNN filtering, GH - GATK HaplotypeCaller with recommended hard filters. ST - Strelka22, FB - Freebayes.

**Supplementary Tables**

**Table S1**. Precision, recall, and F1 statistics of each variant caller for each sample, data type, and variant type (SNP/indel). Table is available as a Supplementary File.

**Table S2**. Comparison of the performance statistics of the pipeline based on BWA MEM and DeepVariant on WES datasets using GRCh38 reference genome with or without decoy sequences.

| Sample | Reference | Variant class | Precision | Recall | F1 Score |
|---|---|---|---|---|---|
| HG002 | Hg38d1 + decoy | SNP | 0.940495 | 0.987303 | 0.963331 |
| | | INDEL | 0.823562 | 0.952271 | 0.883253 |
| | GRCh38.p13 primary | SNP | 0.940123 | 0.987251 | 0.963111 |
| | | INDEL | 0.823331 | 0.952271 | 0.88312 |
| HG003 | Hg38d1 + decoy | SNP | 0.936746 | 0.986257 | 0.960864 |
| | | INDEL | 0.795122 | 0.947715 | 0.864738 |
| | GRCh38.p13 primary | SNP | 0.935753 | 0.986206 | 0.960317 |
| | | INDEL | 0.794894 | 0.947715 | 0.864603 |
| HG004 | Hg38d1 + decoy | SNP | 0.937937 | 0.985502 | 0.961131 |
| | | INDEL | 0.810256 | 0.95251 | 0.875643 |
| | GRCh38.p13 primary | SNP | 0.937511 | 0.985682 | 0.960993 |
| | | INDEL | 0.810256 | 0.95251 | 0.875643 |

Values were calculated using a set of protein-coding and flanking regions (150 bp). Performance was evaluated using the GIAB v. 4.2.1 truth sets.
